# Mapping and characterization of a novel powdery mildew resistance locus (PM2) in *Cannabis sativa* L

**DOI:** 10.1101/2024.12.09.627623

**Authors:** Soren Seifi, Keegan M. Leckie, Ingrid Giles, Taylor O’Brien, John O. MacKenzie, Marco Todesco, Loren H. Rieseberg, Gregory J. Baute, Jose M. Celedon

## Abstract

Breeding genetic resistance to economically important crop diseases is the most sustainable strategy for disease management and enhancing agricultural and horticultural productivity, particularly where the application of synthetic pesticides is prohibited. Powdery mildew disease, caused by the biotrophic fungal pathogen *Golovinomyces ambrosiae*, is one of the most prevalent threats to the cannabis and hemp industry worldwide. In this study, we used bulk-segregant analysis (BSA) combined with high-throughput sequencing to identify and map a novel single dominant resistance (R) locus (designated *PM2*), that strongly suppresses powdery mildew infection and sporulation in *Cannabis sativa*. Histochemical analysis revealed that PM2-induced resistance is mediated by a highly localized hypersensitive response mainly in the epidermal cells of the host. Importantly, genetic markers capable of tracking PM2 resistance in breeding populations were developed using associated SNPs identified in this study.

## Introduction

*Cannabis sativa* L., commonly known as cannabis (marijuana or hemp) is a dioecious, diploid (2n=20), annual, flowering plant species belonging to the Cannabaceae family cultivated for its seed, oil, fiber, and bioactive compounds including cannabidiolic acid (CBDA) and tetrahydrocannabinolic acid (THCA) which have medicinal and psychoactive properties (Pate, 1983; Chandra et al., 2017; Radwan et al., 2017; Kumar et al., 2021). Following the legalization of medicinal and recreational uses of cannabis in many countries in the last few years, cannabis cultivation and related industries have seen a significant expansion (López-Ruiz et al., 2022). Plant diseases caused by fungal, bacterial, and viral pathogens are among the most important factors that threaten cannabis production (Punja, 2021). Powdery mildew (PM) is a widespread and economically damaging fungal disease affecting many indoor, greenhouse, and field grown cannabis crops around the world, including the cannabis industry in Canada and the USA (Pépin et al., 2021). PM disease in cannabis is mainly caused by the obligate biotrophic fungal pathogen *Golovinomyces ambrosiae*, previously known as *G. cichoracearum* (Pépin et al., 2018; Scott and Punja, 2021; Brochu et al., 2022). PM infection attacks the leaves, stems and flowers of cannabis, restricting photosynthesis and nutrient availability, causing premature leaf drop, poor flower quality and significant yield losses (Mihalyov and Garfinkel, 2021; Scott and Punja, 2021). The infection cycle in PM has three main phases: (1) conidium (spore) germination and epidermal penetration; (2) mycelial network development on the host leaf; and (3) conidia generation (asexual reproduction, also known as conidiation or sporulation) (Hückelhoven, 2005). Under optimal conditions, i.e. high humidity and moderate temperature, and access to susceptible cannabis cultivars, PM can complete its life cycle within 1-2 weeks post inoculation (wpi).

PM control in other crops is achieved by application of chemical fungicides, biological control agents, and agricultural practices; however, use of genetically resistant cultivars has historically been the most sustainable, effective, and economical approach in important crops such as wheat, barley and tomato (AGRIOS, 2005; Jorgensen and Wolfe, 2011; Seifi et al., 2014; Bapela et al., 2023). In countries with legal cannabis markets, use of agrochemicals and biological products are tightly controlled by regulatory agencies, and the few available products often exhibit limited protection (Mihalyov and Garfinkel, 2021; Scott and Punja, 2021). Therefore, developing commercial cultivars with genetic resistance to PM remains a highly valuable and sought-after goal in the cannabis industry (Sirangelo, 2023).

The plant immune system comprises several layers of constitutive and inducible defense mechanisms. Constitutive physical and biochemical barriers make up the outer layer of these defenses and can effectively suppress most pathogens. Nevertheless, those few pathogens that succeed in breaking through the preformed defenses will be dealt with by a complex defense machinery induced by the perception of the invading pathogen. Two main mechanisms are involved in the perception of an invading pathogen and the induction of an effective immune response: (i) recognition of pathogen-associated molecular patterns (PAMPs) by membrane-bound receptors leading to PAMP-triggered immunity (PTI), and (ii) detection of specific pathogen-derived effector proteins leading to effector-triggered immunity (ETI). Disease resistance (R) genes typically encode specific nucleotide-binding site leucine-rich repeat (NBS-LRR) proteins that can interact with and detect pathogen-derived effector proteins to induce an ETI response (Jones and Dangl, 2006). Triggering of ETI causes the activation of defense-associated hormonal pathways, typically salicylic acid (SA), and several downstream genes coding for defense responses, such as the hypersensitive response (HR), production of antimicrobial pathogenesis related proteins (PR proteins), and cell wall fortifications, that ultimately deprive the pathogen of the plant’s nutritional resources. (Kosack and Kanyuka, 2007; Shamrai, 2022). R genes are often present in tandem arrays conferring vertical or qualitative resistance in the progeny through dominant Mendelian modes of inheritance where the effect of the dominant (resistance) allele can mask the effect of the recessive (susceptible) one. Qualitative resistance to PM has been reported in hops (*Humulus lupulus*), which is closely related to cannabis (Henning et al., 2017). The first report of a putative R gene mediated resistance against PM in cannabis (named PM1) suggested its location to be on chromosome 2 (Chr ID: NC_044375.1; GenBank acc. no. GCA_900626175.2) (Mihalyov and Garfinkel, 2021). Additionally, a mutation in the susceptibility (S) gene “Mildew Locus O” (*mlo*-mediated loss of susceptibility) has been reported to induce strong resistance to PM in cannabis (Stack et al., 2023).

Bulk-segregant analysis (BSA) coupled with high throughput sequencing has become a popular method for quantitative trait locus (QTL) mapping and is widely used for mapping disease resistance loci within economically important crops (Takagi et al., 2013; Win et al., 2017; Imerovski et al., 2019; Shen et al., 2019; Liang et al., 2020), including those effected by powdery mildew (Ma et al., 2021; Cao et al., 2021). BSA involves the creation of a bi-parental segregating population where both parents display opposing phenotypes for a quantitative or qualitative trait of interest. Unlike traditional QTL mapping where all individuals in a segregating population are genotyped, BSA involves selecting individuals with opposing values for qualitative traits, or individuals from both tails of the distribution for quantitative traits. Selected individuals are grouped into two bulks, representing the extremes of the target trait. This makes BSA attractive for mapping disease resistance QTLs as bulks can be easily made from resistant and susceptible plants. Genotyping in BSA is limited to the two bulks of plants, drastically reducing sequencing cost. Furthermore, several methods of BSA have been developed to utilize single nucleotide polymorphisms (SNPs) called from RNA-Seq data, referred to as BSR-Seq, which provides the required read depth and gene expression data for a lower cost compared to DNA sequencing (Liu et al., 2012; Hill et al., 2013).

Herein, we report the discovery of a novel single dominant PM resistant locus (PM2) that confers strong resistance to PM disease in cannabis. We present evidence indicating that PM2 acts through the induction of a highly localized ROS accumulation in the epidermal and mesophyll layers of the host leaf tissue resulting in HR, the arrest of the pathogen growth, and suppression of its sporulation. We identified two genotypes containing PM2 within a large cannabis diversity population screened for PM resistance. Using BSR-Seq, we mapped PM2 to chromosome 9 (Chr ID: NC_083609.1, Pink Pepper reference genome) and developed genetic markers that can be used to introgress PM2 resistance into elite cannabis commercial cultivars.

## Material and methods

### Plant material

A large diversity population of 510 genotypes called CanD (Cannabis Diversity), including the parental lines W03, N88, and AC, is part of Aurora’s germplasm and is maintained as mother plants to provided clones for experiments. The W03xAC and N88xAC F_1_ mapping populations were generated form a PM-susceptible (AC) parent and PM-resistant (W03 and N88) parents, all described as female, type-I (THC-dominant) drug-type cannabis. Selfed (S_1_) progeny were generated by applying a foliar spray of silver thiosulfate (Millipore Sigma) at a concentration of 0.02M to clones from W03 and N88 parental lines on days 1 and 8 of a short-day photoperiod treatment to induce male flower formation and self-pollination (Mohan Ram and Sett, 1982).

### PM infection assays, disease index, and plant growth conditions

All CanD genotypes were evaluated for PM susceptibility using a clone assay, with at least 6 replicates per genotype, rooted for 14 days in rockwool cubes. Plantlets were inoculated 3 weeks after cloning. Inoculation of healthy plantlets were carried out through “dusting”, whereby fungal spores from a sporulating infected leaf are transferred by tapping the leaf and depositing fresh spores onto the surface of a non-infected leaf. PM spores were sourced from highly infected plants kept in isolation. Disease evaluation and scoring was performed at 4 wpi. Similar methods of inoculation and disease evaluation were used for seedling infection trials. To mimic a production cycle from cloning to harvest, an adult plant assay was performed over the course of a 12-week period. In all assays, plants were grown at 23°C and 80% relative humidity (RH) for the first 48 h (to ensure high levels of spore germination), and then kept at 70% RH during the rest of the infection trial. Photoperiod was 16 h of light and 8 h of dark for clonal propagation and rooting, vegetative growth during the clone assay, and the first two weeks of the adult plant assay. After 2 weeks of growth under vegetative conditions, photoperiod was changed to 12 h of light and 12 h of dark for 10 weeks to induce flowering in the adult plant assay.

Disease severity was assessed with a “disease index” following a method previously developed (Seifi et al., 2013; Seifi et al., 2021), using qualitative severity scores from zero to four according to the area covered by colonies with PM sporulation (“0”, healthy leaves with no signs of infection; “1”, less than 25% coverage; “2”, 25-50% coverage; “3”, 50-75% coverage; and “4”, 75-100% coverage). Disease index was calculated for each genotype by scoring the disease severity in the three most infected leaves from the three most infected plants per genotype using the following formula:

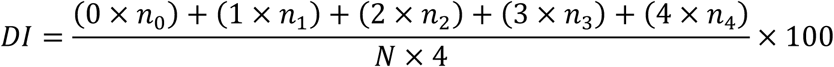

Where DI shows the disease severity in percentage; *n*_0_, *n*_1_, *n*_2_, *n*_3_, *n*_4_ are the number of leaves with disease severity scores of 0, 1, 2, 3 and 4 respectively (Supplemental Figure S1); and N is the total number of leaves evaluated. Disease severity was scored at 4 wpi for clone and seedling assays, and at 12 wpi for adult plant assay.

### Microscopy

The accumulation of hydrogen peroxide was visualized using 3,3-diaminobenzidine (DAB) staining following the protocol devised by Thordal-Christensen et al. (1997). Briefly, samples were treated with DAB-HCl (1 mg/ml) for 3 hours before being fixed in 100% ethanol for downstream microscopy. Fungal structures were stained with 0.1% trypan blue in 10% acetic acid following the protocol described by Seifi et al. (2013). All staining protocols were followed by extensive rinsing steps in demineralized water and samples were subsequently mounted in 50% glycerol before brightfield microscopic observations using an Olympus (Tokyo, Japan) CX443 microscope.

### RNA isolation

W03xAC and N88xAC F_1_ mapping populations were grown from seed in greenhouse conditions under 18 hours of light. At 3 weeks, plants were manually infected with PM by dusting fresh spores as described in the clone infection assay. Evaluations and tissue sampling were conducted at week 7 (4 wpi). For each F_1_ population, equal amounts of leaf samples from 25 resistant and 25 susceptible plants were collected from plants and flash frozen in a slurry of isopropanol and dry ice. Bulks of frozen leaf samples from each population were separately ground in a mortar with liquid nitrogen and 200 mg of ground material was used for RNA isolation using the PureLink^™^ Plant RNA Reagent (ThermoFischer Sci.) following the manufacturer small scale protocol. RNA quality and concentration were assessed using an Agilent Technologies 2100 Bioanalyzer. Only samples with high quality RNA (RNA integrity numbers > 7) were used for RNA sequencing.

### RNA sequencing and SNP calling

Construction of mRNA libraries and sequencing was performed at SBME-Seq center at the School of Biomedical Engineering, University of British Columbia. Each library was sequenced to a depth of ∼20 million PE-reads (150 bp long), using an Illumina NextSeq2000. Library quality and presence of adaptors in raw RNA-Seq reads were analyzed using FastQC (Andrews, 2010). Skewer (Jiang et al., 2014) was used to trim reads to a Phred score no less than Q28. STAR (version 2.7.11a, Dobin et al., 2013) was used to map processed RNA-Seq reads to the Pink Pepper and CBDRx reference genomes. SNP calling was performed by following The Broad Institute’s best practices for RNA-Seq short variant discovery (SNPs + Indels) using GATK (versions 4.3, McKenna et al., 2010). GATK’s Picard implemented MarkDuplicates and SplitNCigarReads tools were used for marking read duplicates and splitting intron spanning reads, respectively. Read duplicates were only marked for aiding variant calling and not removed. GATK’s HaplotypeCaller was used to call variants in each bulk. Resulting variants that did not meet the following quality statistics were removed: quality (QUAL) > 30, quality by depth (QD) > 2.0, Strand Odds Ratio (SOR) < 3.0, FisherStrand (FS) < 60.0, RMS Mapping Quality (MQ) > 40.0, Mapping Quality Rank Sum Test (MQRankSum) > -12.5, and Read Position Rank Sum Test (ReadPosRankSum) > -8.0. Lastly, only bi-allelic (SNPs) variants were selected for BSR-Seq analysis.

### Bulked segregant analysis

BSR-Seq was conducted using a custom in-house R program written to implement both the Euclidean distance metric developed by Hill et al. (2013) and the Bayesian statistical approach developed by Lui et al. (2012). Both methods rely on allele read counts determined by SNP calling to derive allele frequencies used in their calculations. SNPs common to both bulks (susceptible and resistant) were used for analysis. SNPs called from read counts of less than 20 reads were filtered out as their allelic frequencies cannot be accurately measured.

### DNA Isolation

DNA purification was done using lyophilized leaf tissue with a sbeadex Mini Plant DNA Purification Kit^™^ (Biosearch Technologies) on an Oktopure Liquid Handling system.

### Genotyping

Genotyping was done using PCR Allelic Competitive Extension (PACE) 2.0 chemistry (3CR Bioscience, Essex, UK). Assays were designed by the 3CR Bioscience assay design service. All PACE reactions were performed on a QuantStudio 7 (Applied Biosystems) using the manufacturer suggested PACE reaction volumes and cycling conditions. PACE primer sequences are provided in Supplemental Table S1.

## Results

### 1. Germplasm screening for PM resistance

To identify new sources of PM resistance, we screened a total of 510 genotypes in our cannabis germplasm collection, including production cultivars, landraces, and exotic lines, and called this population CanD for **<u>Can</u>**nabis **<u>D</u>**iversity). All 510 genotypes were initially evaluated for resistance/susceptibility to PM using a disease index (DI) measured using a clone infection assay (Materials and Methods). The observed distribution of DI in the CanD population was left-skewed, with more than 70% of the genotypes scoring a DI > 50 indicating a high prevalence of PM susceptibility (Figure 1A). The continuous nature of the distribution of DI in the CanD population suggests that multiple loci, most with small effects, contribute to PM resistance in cannabis.

**Figure 1.**
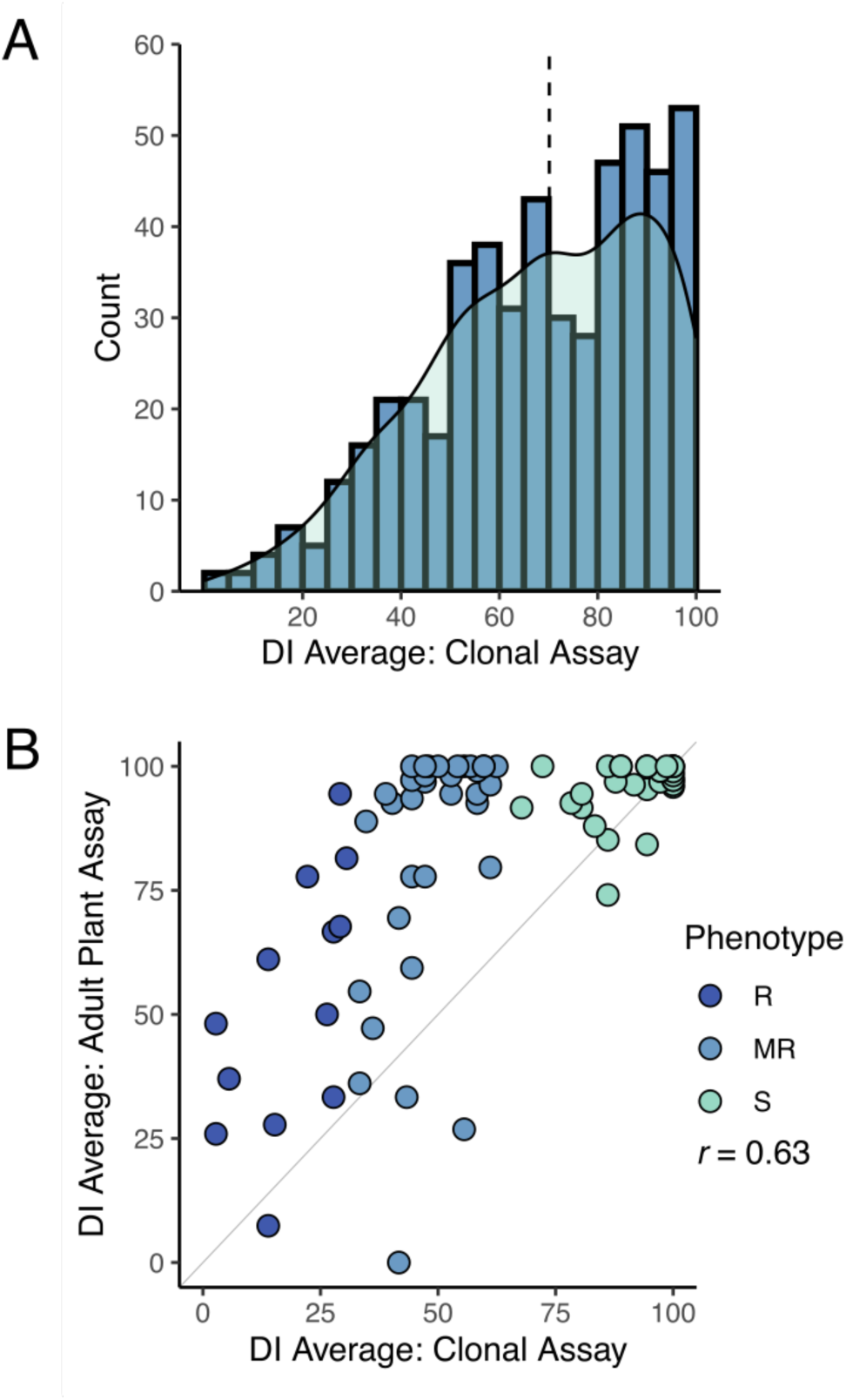
Powdery mildew disease index on cannabis CanD diversity population. (**A**) Distribution of DI results from clonal infection assay. Blue bars indicate number of genotypes within a defined DI range (histogram), while the light green shaded area depicts kernel density estimate of the DI distribution. Black dashed line denotes the median of the distribution. (**B**). DI results from the adult plant infection assay of 90 genotypes selected from the CanD population compared to their corresponding clonal infection assay results. Genotypes from all three resistance categories were selected: Susceptible (S) shown in light green, moderately resistance (MR) shown in blue, and resistant (R) shown in dark blue. The diagonal line (x=y) is shown in light grey as reference. The Pearson correlation between adult infection assay versus clonal infection assay was *r* = 0.63.

To confirm if the resistance to PM observed in clones would persist through the flowering phase of the plant’s life cycle, we performed an adult plant assay on a subset of 90 CanD genotypes. Genotypes were selected to represent the following three phenotypic groups based on their clonal infection assay score: (i) resistant (R: DI ≤ 33), (ii) moderately resistant (MR: 33 < DI < 66), and (iii) susceptible (S: 66 < DI ≤ 100). All resistant genotypes identified in the CanD population were included in the adult plant assay to confirm the resistance to PM observed in clones. Disease pressure in the adult-plant experiment was significantly higher than in the clone assay due to the longer duration (12 vs. 4 weeks) and repeated exposure to high inoculum levels and fresh spores. In the adult plant assay, many genotypes showed an increase in PM susceptibility compared to the clonal assay (Figure 1B, genotypes above the plot diagonal). In severe susceptibility cases, sugar leaves of the flowers, petioles, and parts of the stem tissue were also colonized by the PM pathogen. Overall, the DI in the adult plant assay positively correlated with the results of the clonal assay for the 90 genotypes tested in both experiments (*r* = 0.63). A final group of 12 resistant genotypes (DI < 50) were selected from the adult-plant infection trial. Within this group, 5 genotypes showed strong resistance responses to PM (DI < 33), despite the high disease pressure and length of the assay.

### 2. Mode of inheritance of the observed R locus

Twelve different F_1_ populations derived from highly resistant (R) and susceptible (S) genotypes were subjected to infection trials using the clone assay to determine the mode of inheritance of PM resistance (Materials and Methods). Among the populations tested, two exhibited a 1:1 ratio for R:S in their F_1_ progeny (Figure 2), indicating that resistance to PM in these cases is mediated by a single dominant locus R gene, and that the resistant parents are heterozygous for that locus. The identified resistant parents, W03 and N88 were crossed with a susceptible cultivar, AC (Table 1). The progeny in the remaining nine F_1_ populations tested did not show strong PM resistance, suggesting a multigenic origin of pathogen resistance in the parents of the crosses.

**Figure 2.**
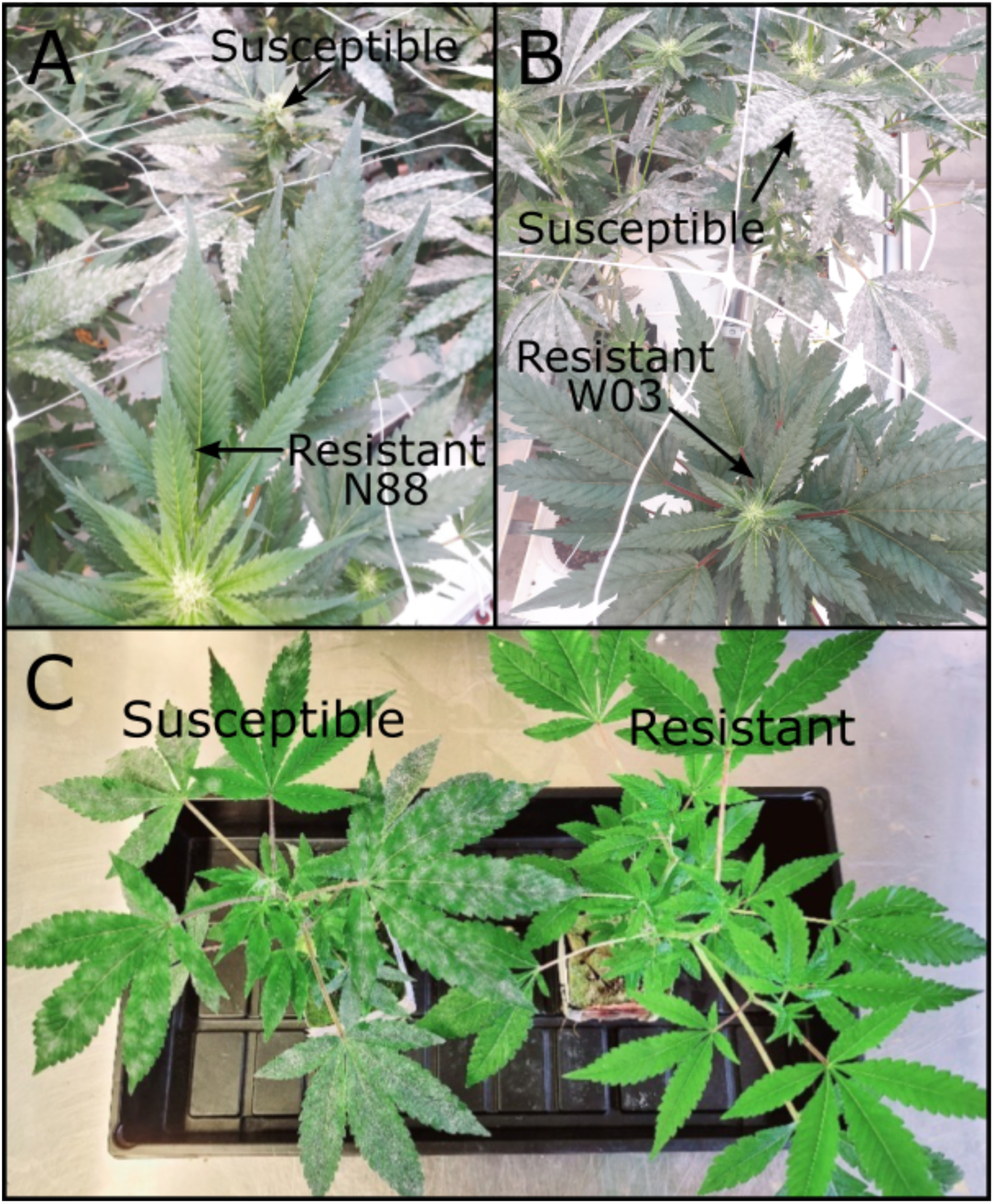
PM resistant genotypes N88 and W03. (**A**) N88 (bottom) next to a PM infected susceptible genotype (top) at 10 wpi during adult plant infection trial. (**B**) W03 (bottom) next to PM infected susceptible genotypes (top) at 10 wpi during adult infection trial. (**C**) Resistant W03xAC-derived F_1_ progeny (right) next to a PM infected susceptible sibling (left) at 4 wpi.

**Table 1:** Chi-squared tests for goodness of fit for the two F_1_ test cross populations.

| Cross (R × S) | Resistant-progeny<br>Exp. vs. Obs. |  | Susceptible-progeny<br>Exp. vs. Obs. |  | Chi-square test |  |  |
| --- | --- | --- | --- | --- | --- | --- | --- |
| | | | | | $\chi^2$ | df | P-value |
| N88 × AC | 140 | 136 | 140 | 138 | 0.007 | 1 | 0.93 |
| W03 × AC | 24 | 24 | 24 | 21 | 0.10 | 1 | 0.90 |

### 3. QTL mapping using Bulk Segregant Analysis

To map the observed dominant PM resistance in the cannabis genome, we performed bulked segregant analysis on the two previously described bi-parental F_1_ populations made from PM-resistant and PM-susceptible genotypes (N88xAC and W03xAC). For each F_1_ population, susceptible and resistant bulks were created consisting of 25 plants each grouped by their respective phenotype (PM susceptible and resistant). RNA sequencing and reference genome alignment was performed on each bulk (Methods and Materials). For the W03xAC population, bulks resulted in 25,150,091 and 17,017,509 reads uniquely aligned to the Pink Pepper Cannabis reference genome for resistant and susceptible bulks respectively and used for SNP calling. A total of 65,202 SNPs shared between both bulks were used for BSR-Seq after filtering for quality scores and read depth (see Methods and Materials). Similarly, the N88xAC population yielded 24,179,503 and 21,576,849 uniquely mapped reads in the resistant and susceptible bulks respectively, and 82,099 shared SNPs after filtering.

To identify the region associated with PM resistance in these populations, we used two separate methods developed for BSR-Seq: the Bayesian implementation by Liu et al. (2012) to estimate the probability of SNPs in linkage disequilibrium with the casual gene, and the Euclidean distance metric developed by Hill et al. (2013), a robust measure of allele frequency differences between bulks. The Bayesian approach identified a cluster of SNPs with high probability of being linked to PM resistance on chromosomes 9 in both the W03xAC and N88xAC populations (Figure 3A). Using the Euclidean distance metric, a strong signal was observed in the same region of chromosome 9 in both populations (Figure 3B). Both the ED and Bayesian probability metrics identify an overlapping region of approximately 2.0Mbp. The boundaries of this region were defined between positions 57,417,178 and 59,418,457 based on SNPs with elevated ED scores and increased posterior probabilities to being linked with PM resistance (Figure 3C-D). To further confirm that the identified region is associated with PM resistance, we repeated the BSR-Seq analysis on a second publicly available reference genome, CBDRx. Using both the Bayesian probability and ED metrics, a cluster of SNPs highly associated with PM resistance was observed on chromosome 9 (NC_044376.1) corresponding to the same region observed in the Pink Pepper reference genome (Supplemental Figure S2). These results strongly suggest that both F_1_ populations are segregating for the same single dominant PM resistance locus, hereafter named PM2.

**Figure 3.**
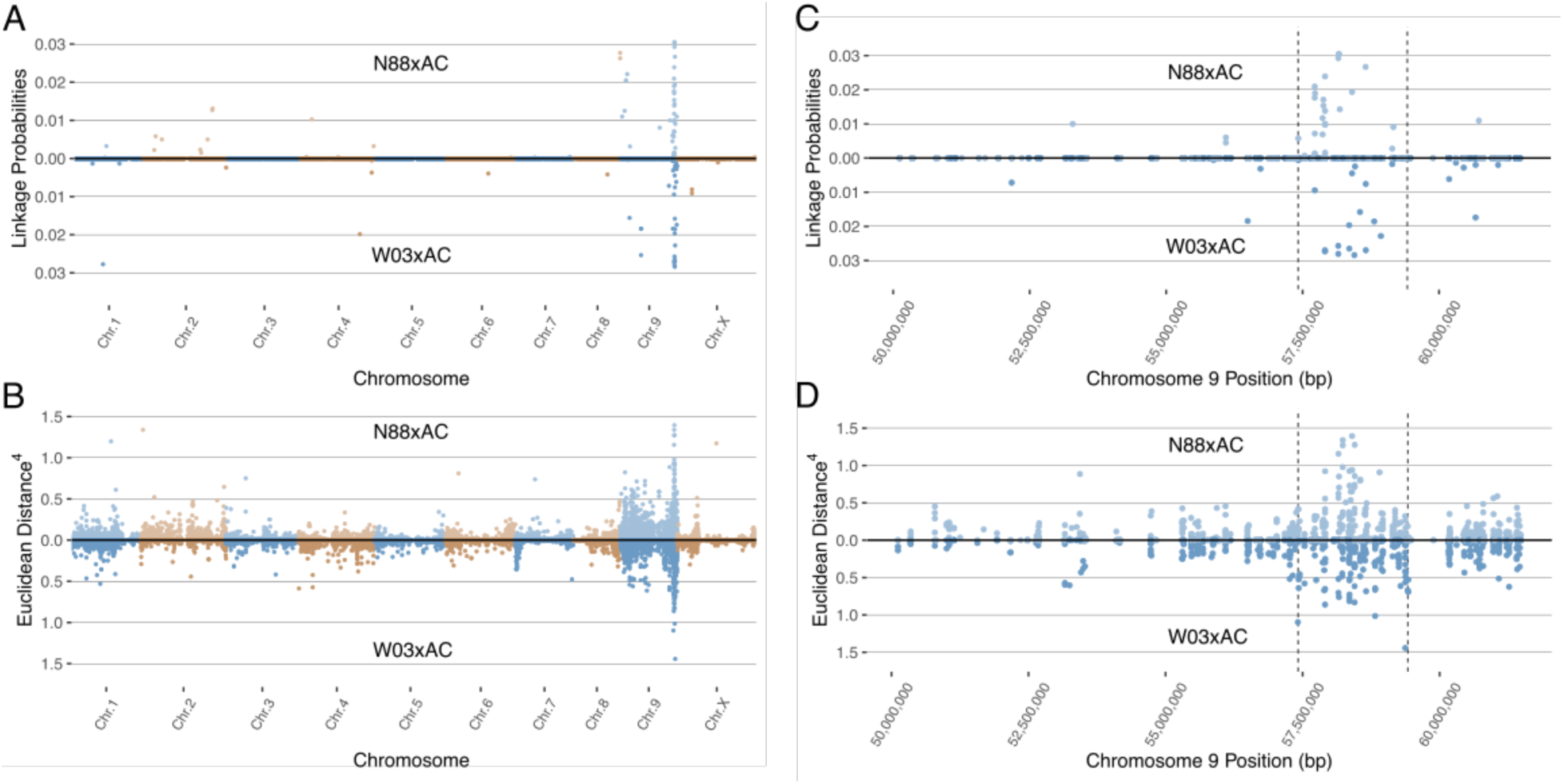
Mapping the PM2 locus using BSR-Seq in both N88xAC (top half) and W03xAC (bottom half) F_1_ segregating populations. Pink Pepper genome was used as reference (GenBank assembly GCA_029168945.1). (**A**) BSR-Seq mapping using the Bayesian implementation to estimate the probability of SNPs linked to PM2 resistance. (**B**) BSR-Seq mapping using the Euclidean distance (ED) metric. ED values for each SNP have been raised to the 4^th^ power to increase signal to noise ratio. Alternating colours denote chromosomes in (**A**) and (**B**). Both Bayesian (**C**) and ED (**D**) metrics identify a region on chromosome 9 between 57,417,178bp and 59,418,457bp containing SNPs associated with PM2 resistance. Vertical dashed lines denote the boundaries of the PM2 associated region, defined by increased Bayesian probability and ED scores.

### 4. Development and Validation of Genetic Markers for Breeding PM2 Resistance

The PM2 QTL defined region contains 2,342 SNPs for the W03xAC population and 2,208 SNPs for the N88xAC population. We developed PACE genotyping assays to validate five SNPs as genetic markers for PM2 resistance in W03xAC and N88xAC F_1_, and W03 and N88 selfed (S_1_) populations (Supplemental Table S2). The accuracy of the markers was tested by comparing their genotype calls to the populations phenotypes as determined by clonal-infection assay. For the W03xAC and N88xAC F_1_ populations, markers had a prediction accuracy between 92 and 99% (Table 2). Marker accuracies were slightly lower when tested on W03 and N88 selfed populations, ranging between 86-93%. The number of genotyped plants in the S_1_ populations were however smaller compared to the number tested in the F_1_ populations (Table 2). To evaluate allelic segregation in the S_1_ populations, a chi-squared test for deviation from Mendelian ratios of inheritance was used and showed that all populations fell within accepted ranges (Table 3). Marker 110 was extensively tested in both F_1_ and S_1_ populations, with its genotype calls and marker accuracy depicted in Figure 4.

**Figure 4.**
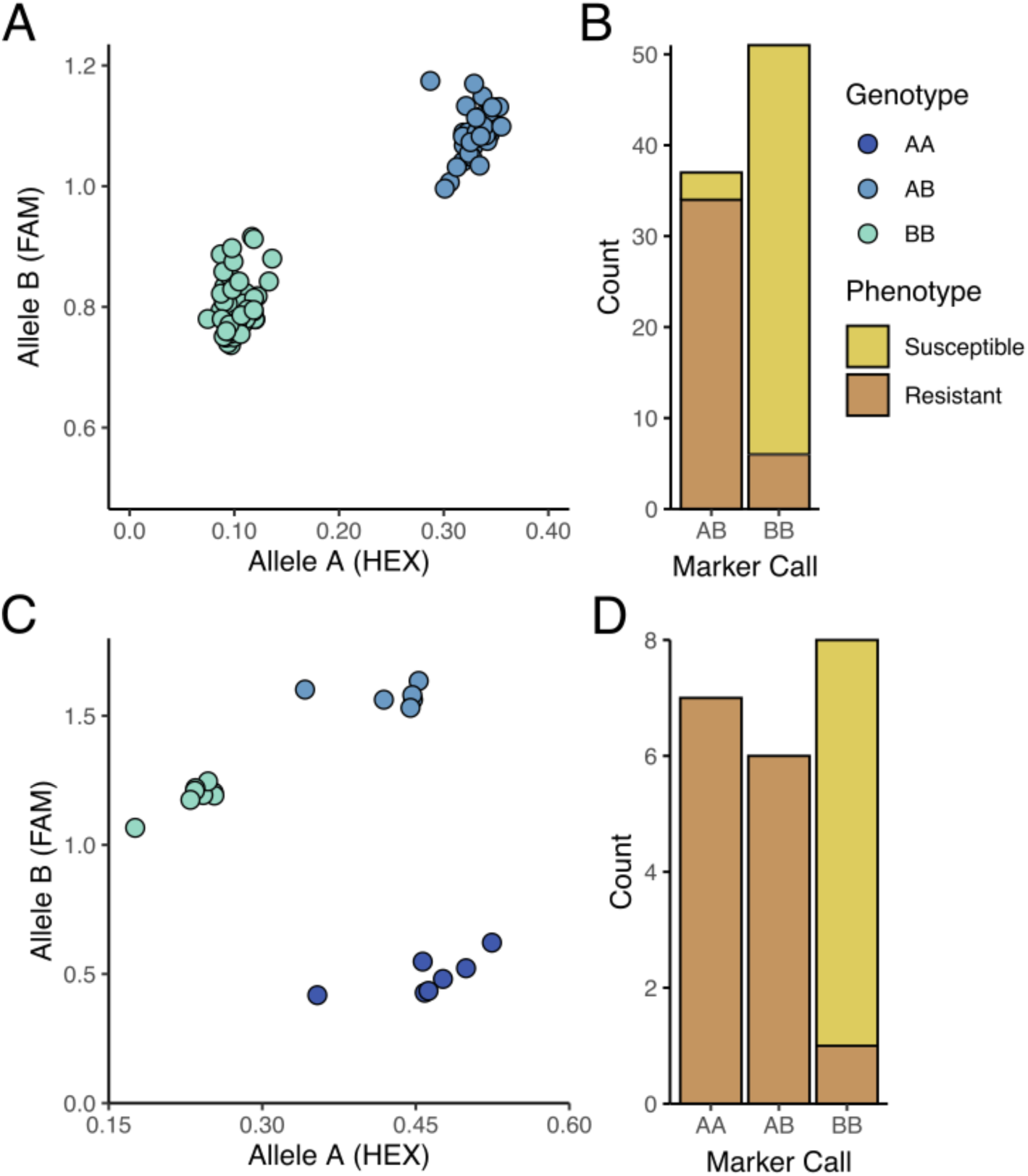
PACE results for marker 110 tracking PM2 mediated resistance. (**A**) Segregation of AB (resistant) and BB (susceptible) alleles in W03xAC F_1_ genotypes. Results depict 1 of 2 independent qPCR runs testing marker 110. (**B**) Accuracy of marker 110 on the F_1_ population from (**A**). Of the 37 F_1_ progeny genotyped as AB, 34 plants were resistant to PM (91.9%). Fifty-one F_1_ progeny were genotyped as BB, with 45 of them susceptible to PM (88.2%). (**C**) Segregation of AA (resistant), AB (resistant), and BB (susceptible) alleles in W03 S_1_ population. (**D**) Accuracy of marker 110 on the S_1_ population from (**C**). Of the 7 S_1_ plants genotyped as AA and 6 S_1_ genotyped as AB, all were resistant to PM. Eight S_1_ plants were genotyped as BB with all but 1 plant being susceptible to PM (87.5%).

**Table 2.**
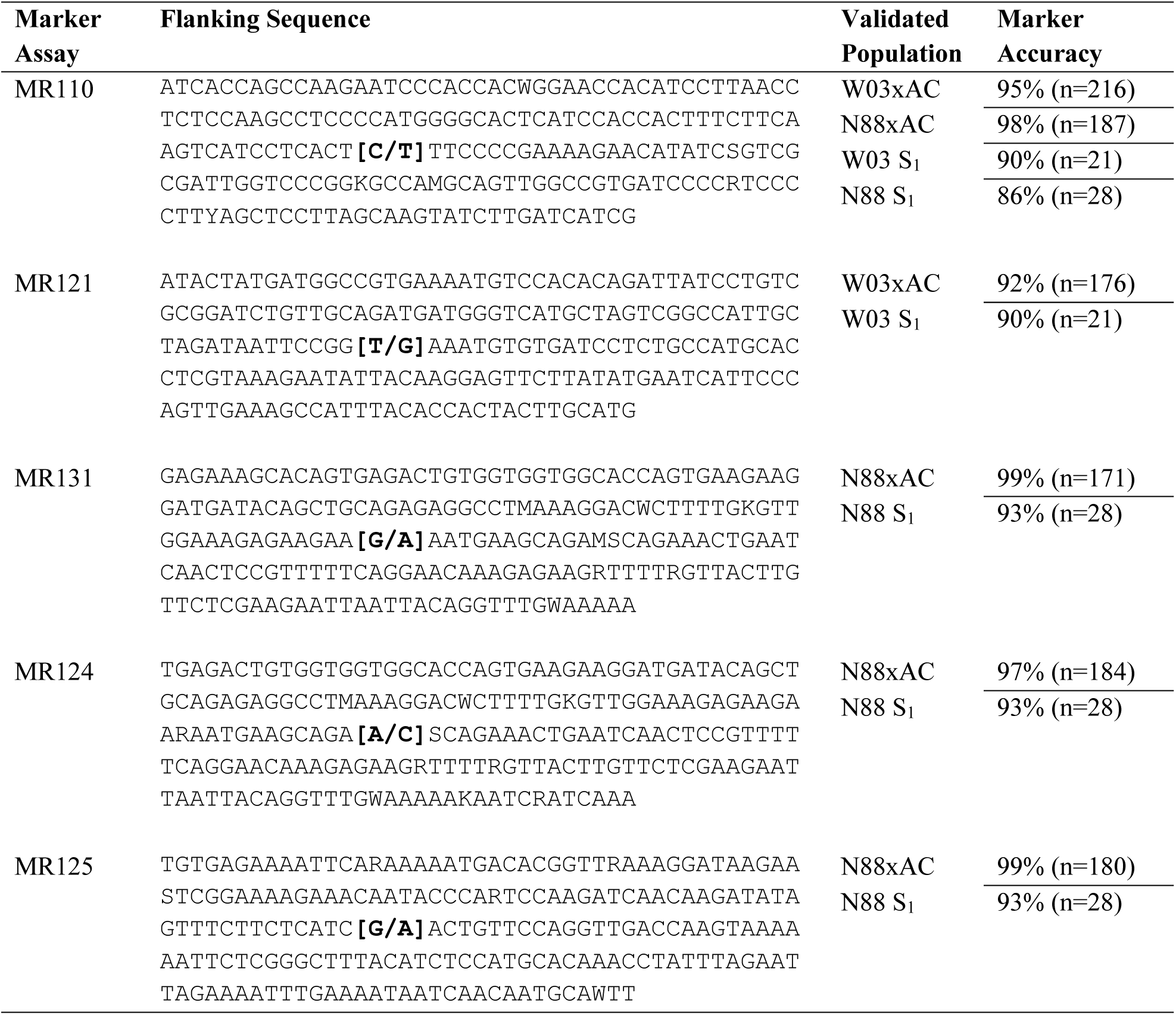
Marker accuracy results and flanking sequence. Marker accuracy for F_1_ populations are the combined results from two independent qPCR runs for each population, resulting in the higher total N value compared to S_1_ populations, which were single runs per populations.

**Table 3.** Chi-squared test for deviation from mendelian patterns of inheritance (df=2).

| Marker | Population | AA | | AB | | BB | | $\chi^2$ | p-value |
| --- | --- | --- | --- | --- | --- | --- | --- | --- | --- |
|  |  | O | E | O | E | O | E |  |  |
| MR 110 | W03 S<sub>1</sub> | 7 | 5.25 | 6 | 10.5 | 8 | 5.25 | 3.9524 | 0.1386 |
| MR 110 | N88 S<sub>1</sub> | 8 | 7 | 12 | 14 | 8 | 7 | 0.5714 | 0.7515 |
| MR 121 | W03 S<sub>1</sub> | 8 | 5.25 | 6 | 10.5 | 7 | 5.25 | 3.9524 | 0.1386 |
| MR 124 | N88 S<sub>1</sub> | 8 | 7 | 11 | 14 | 9 | 7 | 1.3571 | 0.5074 |
| MR 125 | N88 S<sub>1</sub> | 8 | 7 | 12 | 14 | 8 | 7 | 0.5714 | 0.7515 |
| MR 131 | N88 S<sub>1</sub> | 8 | 7 | 11 | 14 | 9 | 7 | 1.3571 | 0.5074 |

Notes: O, observed, E, expected.

### 5. Putative Candidate Genes at the PM2 Locus

Analysis of gene annotations in the PM2 QTL region revealed 13 coding sequences with known functions in disease resistance in other plant species which represent putative candidate genes that could explain PM2 resistance (Figure 5, Table 4). The ratio of expression between resistant and susceptible bulks in some of the selected genes indicated differences that provide additional evidence supporting a putative role in PM2 resistance (Supplemental Table S3).

**Figure 5.**
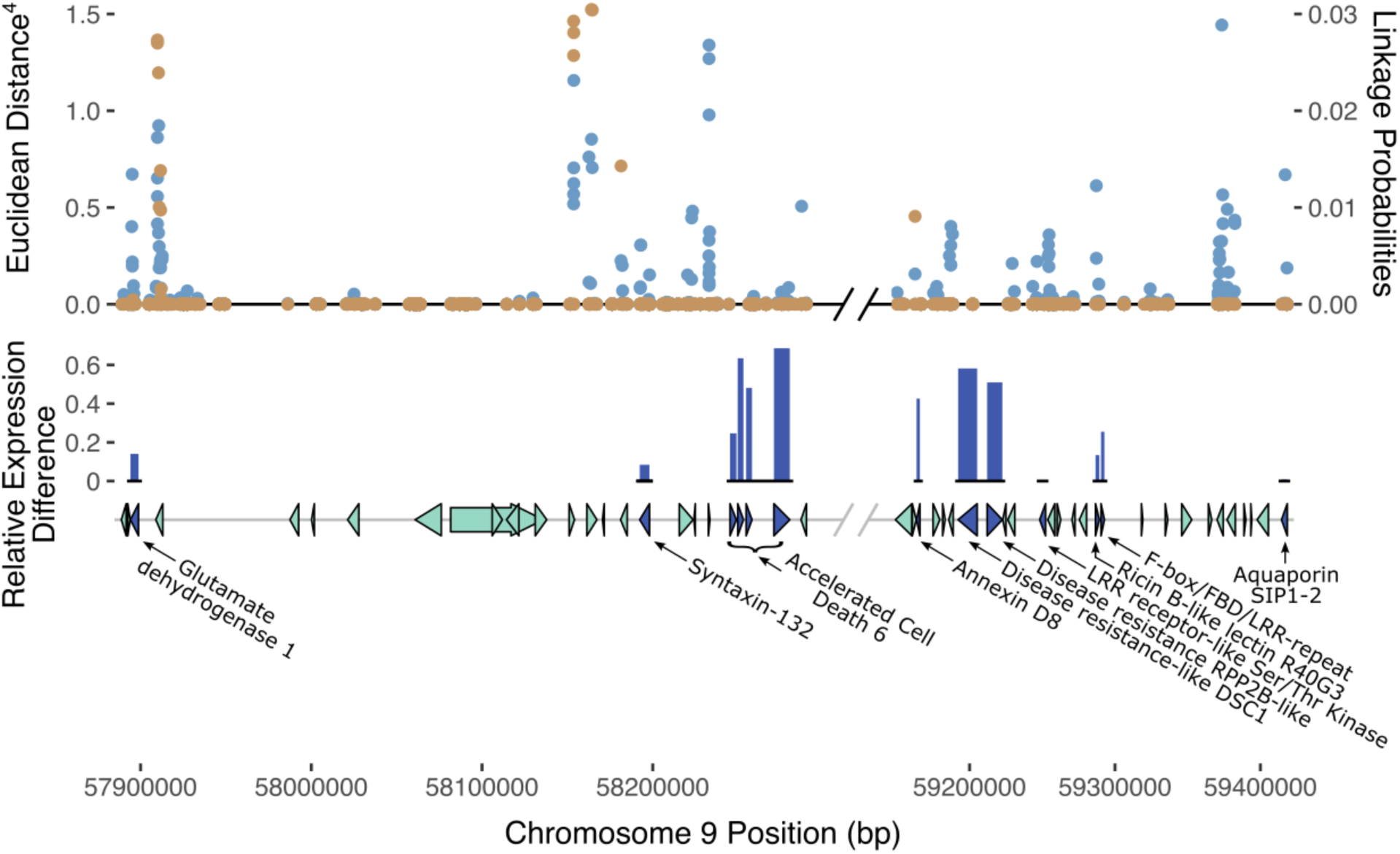
Possible gene candidates mediating PM2 resistance. Gene markers (triangles and arrows) in dark blue denote genes that may be responsible for PM2 resistance based on predicted annotation and previous studies in other plant species. Light green gene markers denote genes less likely to be responsible for PM2 resistance. Non-protein coding genes are not displayed in the gene track. SNPs with their corresponding ED score (left axis) are shown above the gene track in blue (top track). Superimposed in orange are the Bayesian SNP linkage probabilities (right axis). SNP ED and Bayesian probabilities are the combined datasets from both N88xAC and W03xAC F_1_ populations. Height of dark blue bars (middle track) indicate the relative level of gene expression difference between resistant and susceptible bulks for gene candidates. Relative expression difference was calculated as: (Resistance – Susceptible)/Resistance. Expression was measured in transcripts per million (TPM). Diagonal black and grey lines denote a break in chromosome position excluding a region where genes had annotations unrelated to defense responses against pathogens.

**Table 4.**
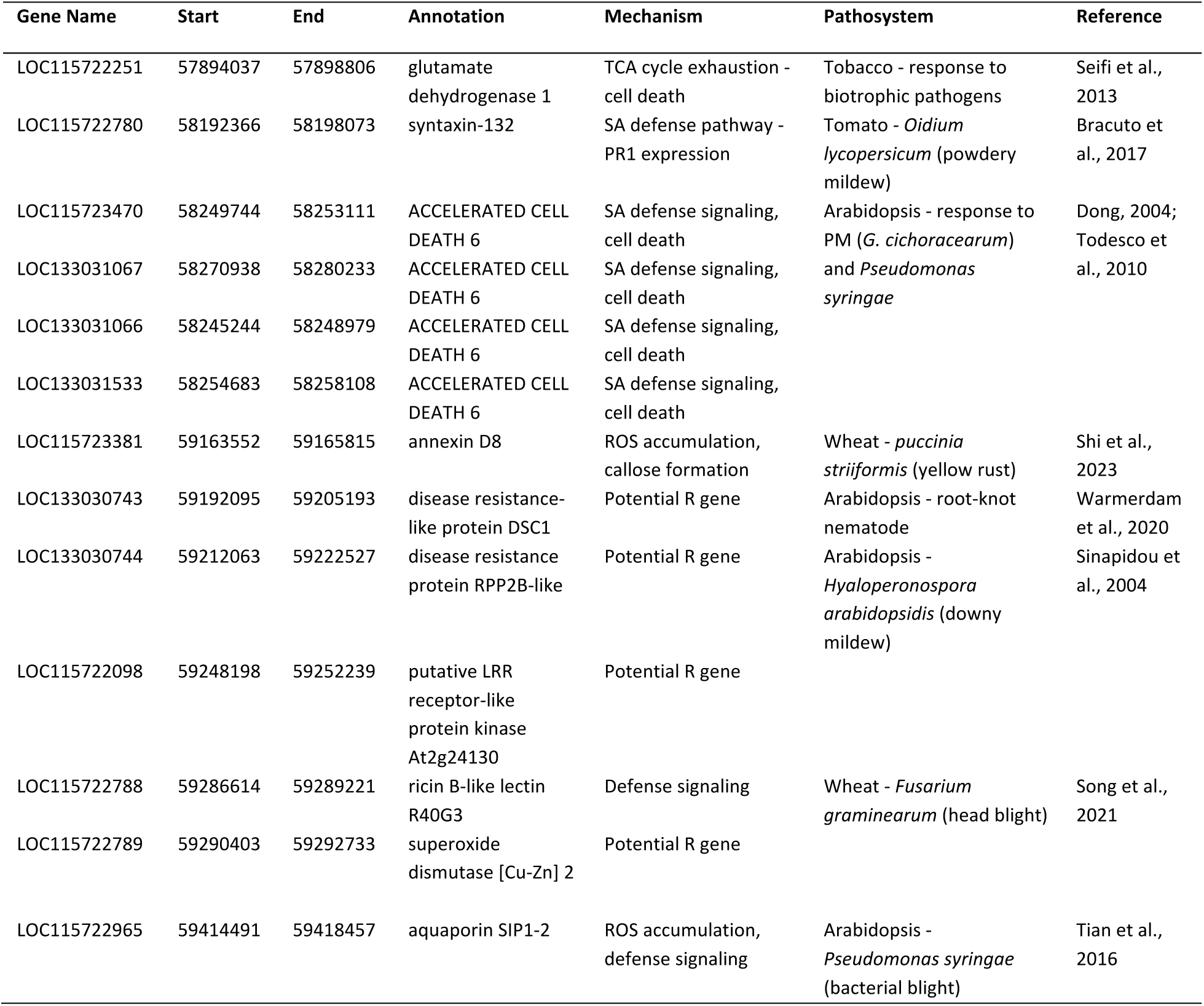
Putative candidate genes within the PM2 region associated with defense signaling and pathogen response. ROS: reactive oxygen species; SA: salicylic acid; PR1: pathogenesis related protein 1.

### 6. Microscopy analysis of PM2-mediated resistance response

To investigate the defense mechanisms underlying PM2 resistance, DAB staining was employed to detect hydrogen peroxide (H_2_O_2_), a key indicator of the hypersensitive response in plants (Seifi et al., 2013). After infecting clones with PM pathogen, PM2-resistant genotypes showed strong and localized DAB staining in the epidermis and mesophyll cells of infected leaves (Figure 6 A-C) while no staining could be observed in the leaves of the susceptible genotype (Figure 6D).

**Figure 6.**
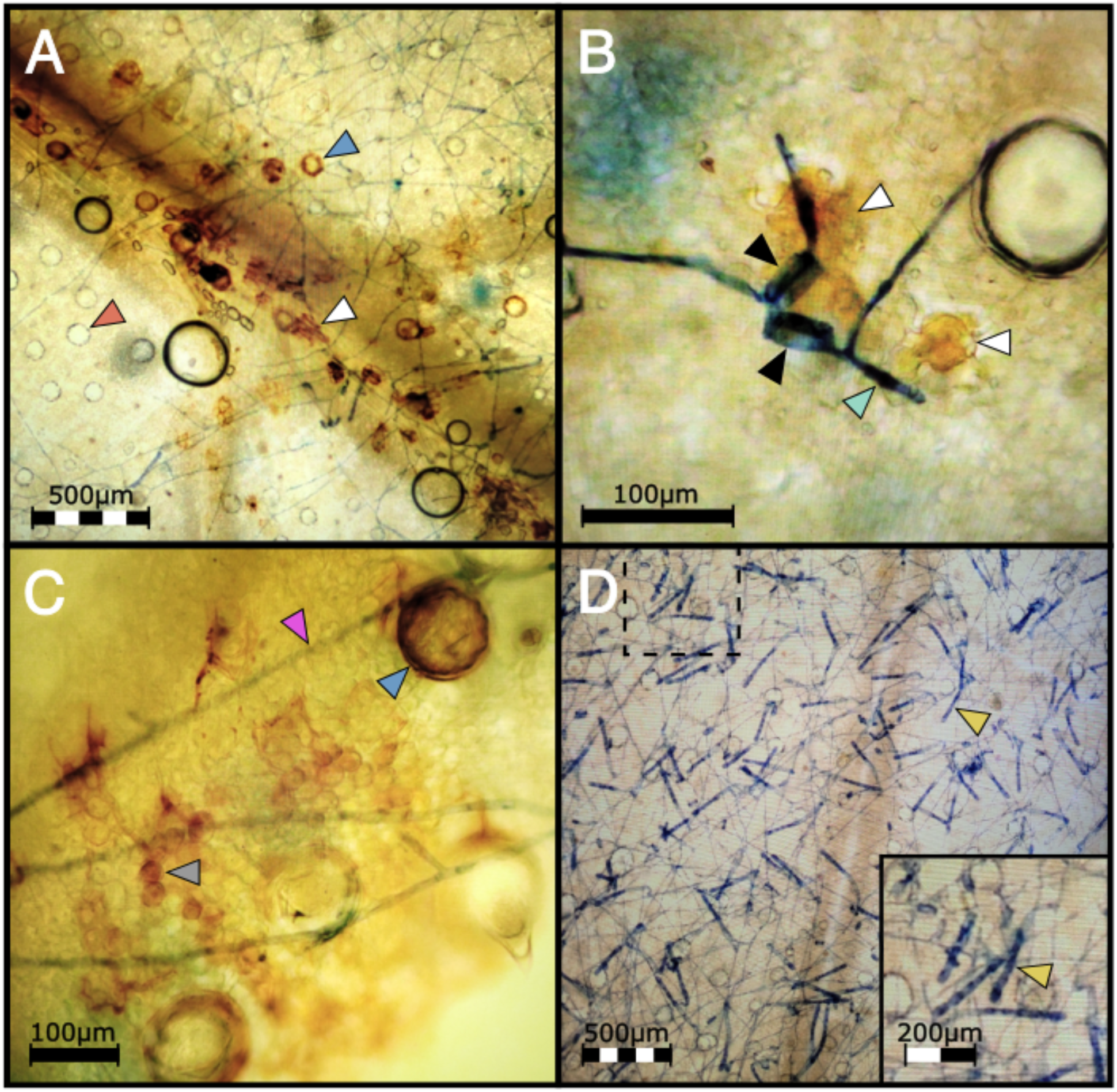
(**A**) DAB staining shows H_2_O_2_ accumulation in a highly localized pattern mainly in epidermal cells (white arrow) in PM2 genotype infected with the mycelia of *G. ambrosiae* (string-like network stained in blue) at 1 wpi. Blue and red arrows show trichome basal cells with and without ROS accumulation respectively. (**B**) Epidermal cells (white arrows) showing ROS accumulation under two germinated PM spores (black arrows) with elongated germ tubes (light green arrow) attempting to penetrate the host tissue. (**C**) Mesophyllic accumulation of H_2_O_2_ (round shape cells, grey arrow) was also observed in PM2 samples under mycelial growth of *G. ambrosiae* (pink arrow). Blue arrow shows a trichome basal cell filled with ROS. (**D**) Susceptible genotype did not show ROS accumulation after DAB staining and showing high levels of conidiophore (yellow arrow) production instead. Dashed box denotes the magnified region in bottom right inset.

In susceptible cannabis genotypes, the PM fungal pathogen *G. ambrosiae* fully develops its mycelial network and conidiophores within 2 wpi, completing its infectious life cycle. Disease development was compared between W03, AC, and a third genotype, P04, having an unknown form of PM resistance (PMU) identified in our clonal infection assay (Figure 7A). Clear differences in infection symptoms, mycelia growth, and pathogen reproduction were observed between PM2-mediated resistance and PMU. PM2-mediated resistance in W03 showed restricted mycelial growth and strong suppression of the conidiophore formation, which is a key developmental stage for the sporulation phase of the pathogen (Figure 7B). In contrast, susceptible genotype AC under identical conditions showed dense mycelial growth with high amounts of conidiophore formation (Figure 7C). While conidiophore formations and sporulation were largely suppressed in the PM2-mediated response, normal conidiophore formation and sporulation occurred in sporadic patches of restricted PM colonies in the PMU-induced response. It should be noted that P04 scored a disease index value of 16.77 in our clonal infection assay and was therefore deemed resistant. The reduced level of infection observed in W03 and N88 compared to P04 highlights the efficacy of PM2-mediated resistance.

**Figure 7.**
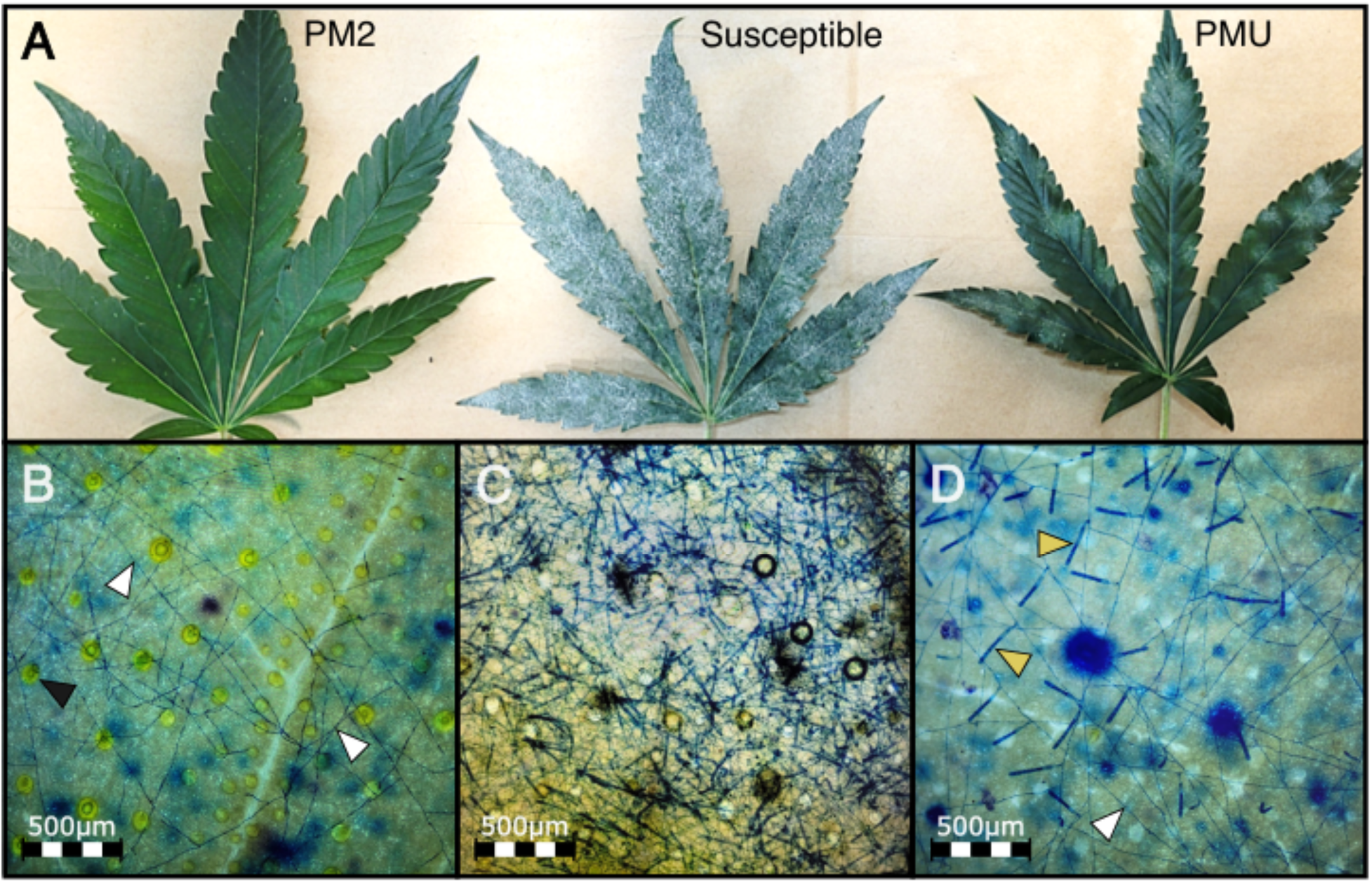
PMU- and PM2-mediated response compared to a susceptible cannabis genotype at 4 wpi. (**A**) PM infected leaves from PM2 carrying genotype (left), susceptible genotype (center) and PMU carrying genotype (right). (**B**) Mycelial network (white arrows) developed on PM2 resistant genotype with no conidiophores. Black arrows denote basal trichome cells. (**C**) High density of conidiophore and sporulation can be seen on the susceptible genotype. (**D**) Low density of conidiophore (yellow arrows) generation can be seen in sporadic patches of PM mycelial growth (white arrow) on PMU.

To validate the repression of conidiophore formation observed in PM2-mediated resistance, number of conidiophores per 3.8 mm^2^ of leaf surface were counted at 9 randomly selected positions on leaves of W03, P04, and susceptible genotype AC (Figure 8). Significant differences in mean conidiophore count were observed in all three genotypes [Kruskal-Wallis test, *X*^2^ (2, 27) = 23.15, *p* < 0.001]. Both mean conidiophore counts from W03 and P04 were significantly lower compared to susceptible AC, *p* < 0.001 and *p* < 0.05 respectively (Dunn’s *post-hoc* test with Bonferroni correction). When comparing PM2-mediated resistance to PMU, W03 had significantly lower counts of conidiophores compared to P04 (*p* < 0.05), with mean number of observed conidiophores at 5.33 compared to 35.44 for P04, and 118.33 for AC. These results further demonstrate the robust level of PM resistance conferred by PM2.

**Figure 8.**
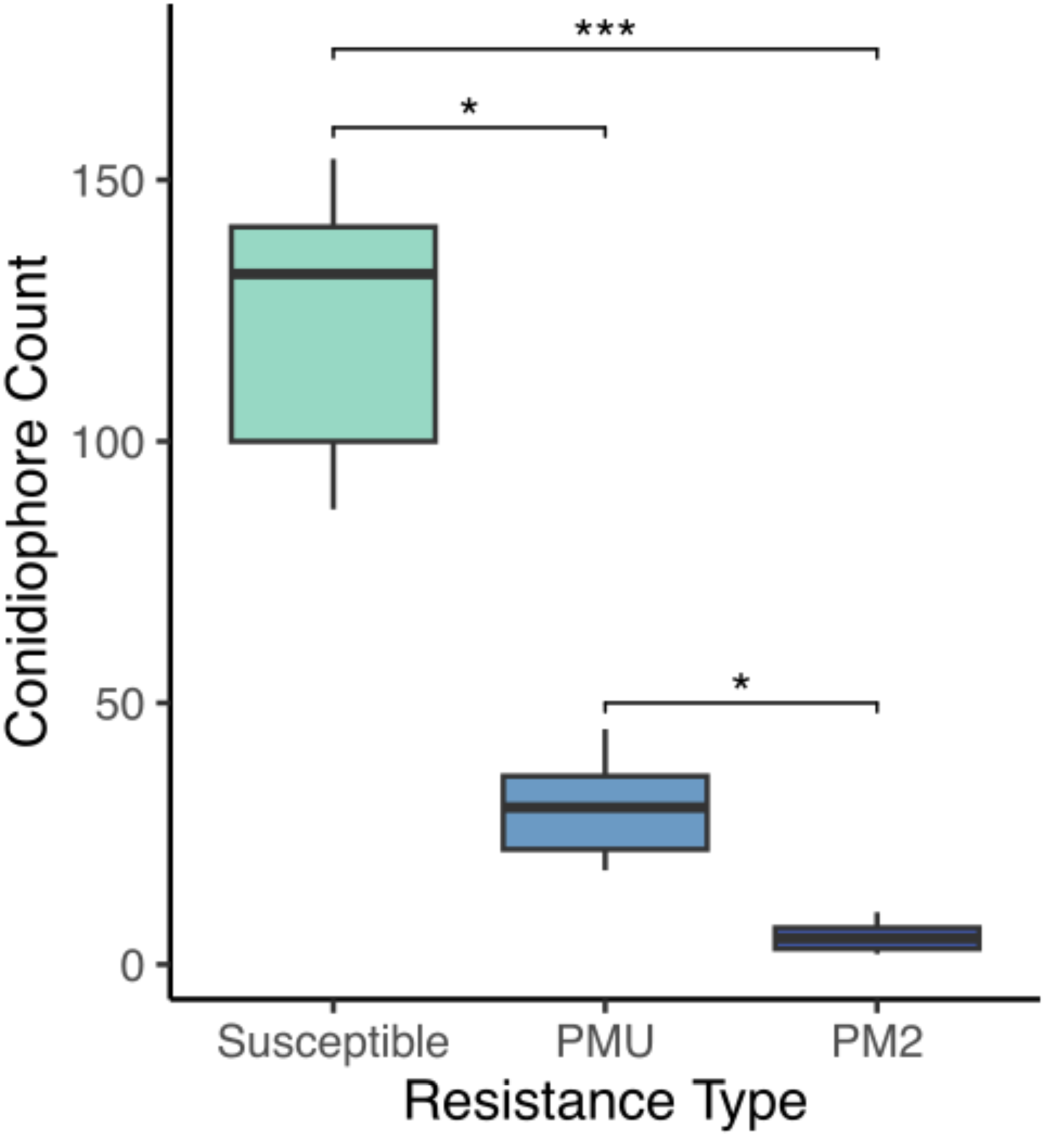
Powdery mildew conidiophore density observed on susceptible genotype AC, PMU resistant genotype P04, and PM2 resistant genotype W03. Values indicate mean number of conidiophores observed per 3.8 square mm of leaf surface with 9 replicates per resistant type. Asterisks denote level of significance between groups [Kruskal-Wallis test, *X*^2^ (2, 27) = 23.15 with Dunn’s post-hoc test and Bonferroni correction (*P*\* < 0.05, *P*\*\*\* < 0.001)].

## Discussion

Powdery mildew (PM) is the most prevalent fungal disease in indoor cannabis growing operations, causing significant losses to the industry (Pépin et al., 2021). Qualitative resistance to PM has been reported in hops (Humulus lupulus), a species closely related to cannabis (Henning et al., 2017). Recently, the first report of a single dominant resistance locus against PM was reported in cannabis, named PM1, with a suggested location of the causal R gene on chromosome 2 (CS10 Chr ID: NC_044375.1; GenBank acc. no. GCA_900626175.2) (Mihalyov and Garfinkel, 2021). In this study, we identified a new single dominant PM resistance locus, named PM2, located on chromosome 9 (CS10cChr ID: NC_044376.1). No macroscopic chlorotic spots were detected on PM2 leaves. However, DAB staining revealed H_2_O_2_ accumulation in epidermal and mesophyll cells beneath the pathogen’s mycelial growth, indicating a highly localized HR reaction in PM2-mediated resistance. Furthermore, the reproduction phase of the fungal pathogen was strongly suppressed, resulting in a significant reduction of conidia generation (>90%) in PM2 genotypes. This could be explained by the inhibition of penetration through HR by the resistant genotype, leading to the observed long ‘wandering’ mycelial growth with delayed and minimal reproduction. Such a phenomenon has been previously reported in an interaction between the biotrophic rust pathogen *Puccinia striiformis* and a resistant cultivar of wheat harboring the YR15 R gene (Seifi et al., 2021).

Further exploration focused on defense-associated genes flanking PM2 revealed several annotated genes (Table 5). Notably, these genes fall into three main functional categories: 1) hormonal regulation, particularly SA signaling pathway; 2) genes involved in ROS accumulation and cell death induction; and 3) genes predicted to encode potential R proteins, including one (LOC115722098) exhibiting LLR and Ser/Thr-kinase domains. Studies in other plant species have highlighted the role of annexin and aquaporin proteins in defense responses against biotrophic pathogens. For example, annexin proteins regulate SA-dependent defense responses and influence ROS generation and callose deposition (Shi et al., 2023), while aquaporin proteins link extracellular ROS accumulation to intracellular defense signaling in Arabidopsis challenged by pathogens (Tian et al., 2016). *RPP2B*, an R gene with a TIR-NB-LRR domain, plays a crucial role in race-specific recognition and defense signaling against downy mildew (Sinapidou et al., 2004). Similarly, the TIR-NB-LRR DSC1 protein is implicated in basal immunity against root-knot nematodes in Arabidopsis (Warmerdam et al., 2020). Research on the TCA cycle through glutamate dehydrogenase suggests its role in defense responses against biotrophic pathogens, including the induction of programmed cell death (Seifi et al., 2013). Accelerated Cell Death 6 (ACD6), involved in SA-mediated defense responses in Arabidopsis, triggers programmed cell death and expression of PR genes against bacterial pathogens (Dong et al., 2004; Chika et al., 2014). Notably, ACD6 belongs to the transmembrane-ankyrin-repeat protein family with a key regulating role in tradeoffs between vegetative growth and general pathogen defense, conferring broad-spectrum disease resistance to pathogens including PM in Arabidopsis (Todesco et al., 2010). Syntaxin proteins have been identified as crucial players in plant defense responses to PM. Studies on barley and Arabidopsis suggest that syntaxin genes regulate fungal penetration and enhance resistance against PM pathogens (Bracuto et al., 2017). Notably, silencing of the syntaxin gene SlPEN compromised resistance in a *mlo*-resistant tomato line against *Oidium neolycopersici*, highlighting their importance in PM resistance (Bracuto et al., 2017).

Originally developed using restriction fragment length polymorphisms (RFLPs) and random amplified polymorphic DNAs (RAPDs) markers (Michelmore et al., 1991), BSA now employs the use of next-generation sequencing (NGS) technologies which have greatly improved its power of detection in mapping phenotypic traits. Algorithms designed for detecting marker associations to target traits have been developed for NGS-based BSA (also referred to as BSA-Seq). These algorithms are primarily based on detecting differences in allele frequencies between bulks, paired with smoothing techniques that help identify positive signals from noise. Popular BSA-Seq statistics include the G’ value (Magwene et al., 2011), SNP-index/InDel-index (Takagi et al., 2013; Singh et al., 2017), Euclidean distance (ED, Hill et al., 2013), and smoothLOD (Zhang et al., 2019), among others (see Li et al., 2022). In addition to whole genome resequencing, bulk segregant analysis using SNP markers called from transcriptome data (BSR-Seq) has been successfully used to map QTLs in maize (Liu et al., 2012), zebrafish (Hill et al., 2013), pacific white shrimp (Dai et al., 2018), pea (Wu et al., 2022) and wheat (Trick et al., 2012; Li et al., 2018; Hao et al., 2019). Several BSA statistics have been developed specifically to handle the challenges inherent to RNA-Seq data, such as allele-specific expression and variable coverage. These include a Bayesian approach to estimate marker association (Liu et al., 2012) and the ED metric (Hill et al., 2013), both of which were successfully used in the present study to map PM2 mediated resistance in cannabis. Both the Bayesian and Euclidean distance methods identified the same 2Mb region within chromosome 9, containing the PM2 QTL. While the Bayesian method required considerably longer computational times, the signal to noise ratio was superior to ED, leaving an easily identified signal peak associated with PM resistance. The comparison of two mapping populations segregating for PM2 provided further strength to our analysis. Both populations showed clear signals for PM2 mediated resistance in the same location on chromosome 9. A minor secondary peak was observed on chromosome 7, however this was not investigated in this study as it was observed in the W03xAC F_1_ population only.

The use of RNA-Seq data for SNP calling provides the added benefit of gene expression data from both resistant and susceptible bulks. Several gene candidates for PM2 mediated resistance identified by BSR-Seq showed increased gene expression in resistant bulks compared to susceptible bulks, including *ACD6*, *Annexin D8*, *RPP2B*, and *DSC1*. This gene expression data, however, comes from pooled samples with no biological replicates, and therefore should not be used to draw conclusions, but to guide follow-up studies.

Populations frequently used in BSA include bi-parental populations segregating for a trait of interest. Common types include F_2_, F_2:3_, BC_1_, and recombinant inbred lines (Zou et al., 2016). However, any population containing individuals with contrasting traits can in theory be used in a bulked sample analysis (Li et al., 2022). For species that have long life cycles, or are not amenable to selfing, F_1_ populations have been used (Shen et al., 2019; Dougherty et al., 2018). These populations require parents to be highly heterozygous and have observable phenotypic variation in the F_1_ generation, as is the case with dominant traits (Li et al., 2022). Both our N88xAC and W03xAC F_1_ populations exhibited clear segregation in PM resistance; employing these populations saved a considerable amount of time, by removing the need to proceed to the F_2_ generation.

The development of resistant cultivars is an effective practice for the management of PM disease control. Successful introgression of PM2 mediated resistance into elite cannabis cultivars will reduce the reliance of chemical pesticides, which are heavily regulated in cannabis as in most other crops. As the cannabis industry expands by developing new markets and applications, there will be an increased need for resistant cannabis cultivars, highlighting the relevance of studies, such as the one presented here, identifying the genetic basis of resistance and susceptibility to pathogens in this species.

## Supporting information

Supplementary Material Seifi et al.

## Acknowledgments

We would like to thank Dr. Pauline Duriez for her valuable support and helpful discussions in this study.

## Data Availability Statement

RNA-Seq data used for BSR-Seq in both N88xAC and W03xAC F_1_ populations has been uploaded to NCBI’s sequence archive under BioProject ID PRJNA1182366.

