## Supplementary Material Seifi et al. for "Mapping and characterization of a novel powdery mildew resistance locus (PM2) in *Cannabis sativa* L"

**Supplemental Material**

**Supplemental Figure S1**. Illustration of the severity scale of PM infection on cannabis leaves used for disease index evaluation.

**Supplemental Figure S2.** Mapping the PM2 locus using BSA in the CS10 CBDRx reference genome.

**Supplemental Table S1.** PACE Sequence primers used for genetic markers tracking PM2 resistance.

**Supplemental Table S2.** Genetic markers associated with PM2 resistance and their location in reference genomes.

**Supplemental Table S3.** RNA-Seq gene expression data for genes within the PM2 locus for both W03xAC and N88xAC F1 bulks.

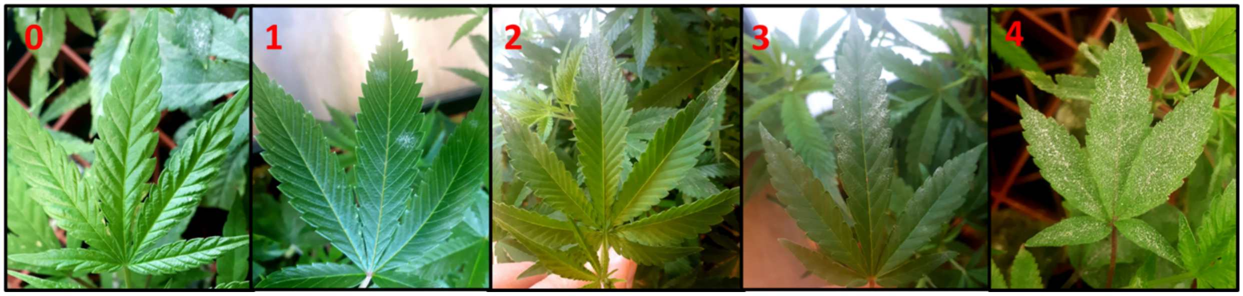


**Supplementary Figure S1.** Illustration of the severity scale of PM infection on cannabis leaves used for disease index evaluation. **0**: no symptoms; **1**: few restricted (not spreading) colonies; **2**: PM colonizing less than 50% of the leaf; **3**: PM colonizing more than 50% but not the whole leaf; **4**: PM colonizing the whole leaf surface.

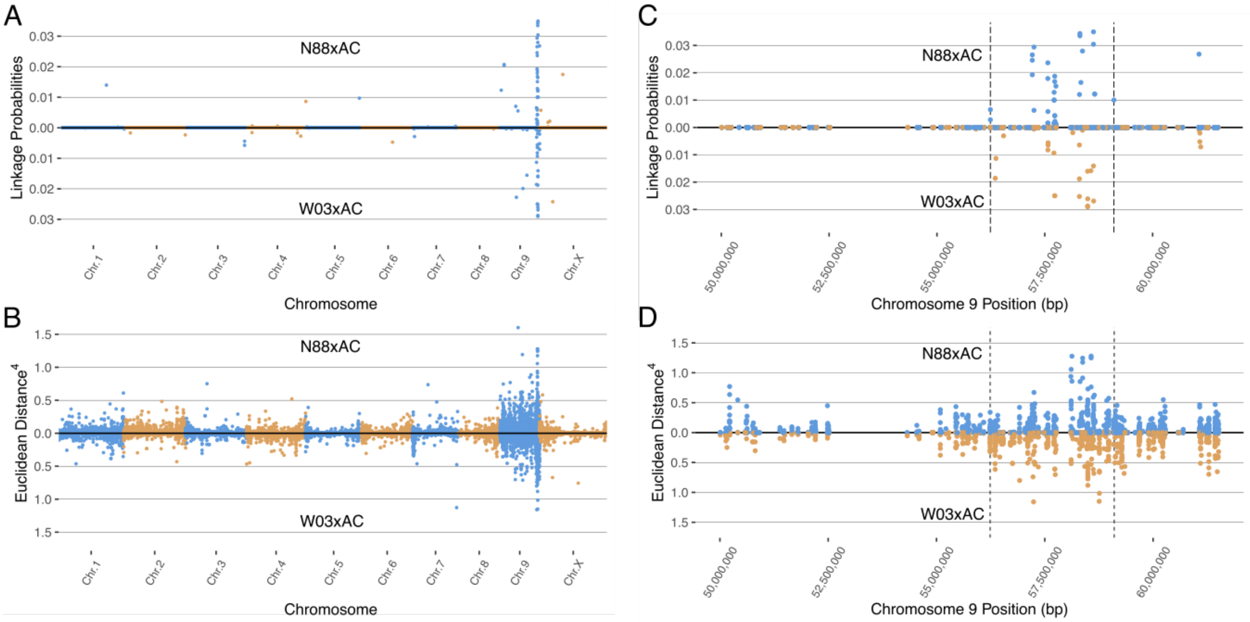


**Supplemental Figure S2.** Mapping the PM2 locus using BSA in both N88xAC (top half) and W03xAC (bottom half) F1 segregating populations. CBDRx (GenBank assembly GCA_900626175.2) genome was used as reference. **(A)** BSA mapping using the Bayesian implementation to estimate the probability of SNPs linked to PM2 resistance. **(B)** BSA mapping using the Euclidean distance (ED) metric. ED values for each SNP have been raised to the 4$^{th}$ power to increase signal to noise ratio. Alternating colours denote chromosomes in **(A)** and **(B)**. Both Bayesian **(C)** and ED **(D)** metrics identify a region on chromosome 9 between 56,237,869 and 59,102,577 containing SNPs associated with PM2 resistance. Vertical dashed lines denote the boundaries of the PM2 associated region, defined by increased ED scores. Blue colour denotes values for N88xAC while orange denotes values for W03xAC in **(C)** and **(D)**.

**Supplemental Table S1**. PACE sequence primers used for genetic markers tracking PM2 resistance.

| Marker | Forward Primer 1 | Forward Primer 2 | Common Reverse Primer |
|---|---|---|---|
| MR110 | CACTTTCTTCAAGTCATCCTCACTC | CCACTTTCTTCAAGTCATCCTCACTT | CGACSGATATGTTCTTTTCGGGGAA |
| MR121 | GGCCATTGCTAGATAATTCCGGT | GGCCATTGCTAGATAATTCCGGG | GTGCATGGCAGAGGATCACACATTT |
| MR124 | TGGAAAGAGAAGAARAATGAAGCAGAA | GGAAAGAGAAGAARAATGAAGCAGAC | GTTCCTGAAAAACGGAGTTGATTCAGTTT |
| MR125 | TTACTTGGTCAACCTGGAACAGTC | TTTACTTGGTCAACCTGGAACAGTT | ACCCARTCCAAGATCAACAAGATATAGTTT |
| MR131 | TTTGKGTTGGAAAGAGAAGAAG | CTTTTGKGTTGGAAAGAGAAGAAA | GTTCCTGAAAAACGGAGTTGATTCAGTTT |

**Supplemental Table S2**. Genetic markers associated with PM2 resistance and their location in reference genomes.

| Marker | Ref | Alt | Pink Pepper | | CBDRx | |
|---|---|---|---|---|---|---|
| | | | Chromosome | Position | Chromosome | Position |
| MR110 | C | T | NC_083609.1 | 58,448,827 | NC_044376.1 | 58,108,802 |
| MR121 | G | T | NC_083609.1 | 58,552,298 | NC_044376.1 | 58,570,897 |
| MR124 | A | C | NC_083609.1 | 58,289,703 | NC_044376.1 | 58,423,201 |
| MR125 | G | A | NC_083609.1 | 58,164,119 | NC_044376.1 | 58,314,046 |
| MR131 | G | A | NC_083609.1 | 58,289,715 | NC_044376.1 | 58,423,213 |

Notes: Chromosome coordinates of markers are provided in both Pink Pepper and CBDRx reference genomes. Ref indicates reference allele found in the Pink Pepper reference genome and Alt the alternative allele.

**Supplemental Table S3.** Provided as a standalone excel file “Supplemental Table S3.xlsx”.
